# Identification of *Slco1a6* as a candidate gene that broadly affects gene expression in mouse pancreatic islets

**DOI:** 10.1101/020974

**Authors:** Jianan Tian, Mark P. Keller, Angie T. Oler, Mary E. Rabaglia, Kathryn L. Schueler, Donald S. Stapleton, Aimee Teo Broman, Wen Zhao, Christina Kendziorski, Brian S. Yandell, Bruno Hagenbuch, Karl W. Broman, Alan D. Attie

## Abstract

We surveyed gene expression in six tissues in an F_2_ intercross between mouse strains C57BL/6J (abbreviated B6) and BTBR *T*^+^ *tf* /J (abbreviated BTBR) made genetically obese with the *Leptin*^*ob*^ mutation. We identified a number of expression quantitative trait loci (eQTL) affecting the expression of numerous genes distal to the locus, called *trans*-eQTL hotspots. Some of these *trans*-eQTL hotspots showed effects in multiple tissues, whereas some were specific to a single tissue. An unusually large number of transcripts (∼8% of genes) mapped in *trans* to a hotspot on chromosome 6, specifically in pancreatic islets. By considering the first two principal components of the expression of genes mapping to this region, we were able to convert the multivariate phenotype into a simple Mendelian trait. Fine-mapping the locus by traditional methods reduced the QTL interval to a 298 kb region containing only three genes, including *Slco1a6*, one member of a large family of organic anion transporters. Direct genomic sequencing of all *Slco1a6* exons identified a non-synonymous coding SNP that converts a highly conserved proline residue at amino acid position 564 to serine. Molecular modeling suggests that Pro564 faces an aqueous pore within this 12-transmembrane domain-spanning protein. When transiently overexpressed in HEK293 cells, BTBR OATP1A6-mediated cellular uptake of the bile acid taurocholic acid (TCA) was enhanced compared to B6 OATP1A6. Our results suggest that genetic variation in *Slco1a6* leads to altered transport of TCA (and potentially other bile acids) by pancreatic islets, resulting in broad gene regulation.

## Introduction

The measurement of genome-wide gene expression levels in segregating populations (such as the offspring of a cross between two inbred mouse strains), an endeavor termed genetical genomics (Jansen and Nap 2001) or expression genetics (Broman 2005), offers the promise of accelerating the identification of genes contributing to variation in complex phenotypes and further enables the genetic dissection of gene expression regulation (reviewed in Albert and Kruglyak 2015). Early experiments in yeast (Brem *et al.* 2002; Yvert *et al.* 2003), mouse (Schadt *et al.* 2003), and human (Cheung *et al.* 2003; Morley *et al.* 2004) revealed some of the basic features of the genetic architecture of gene expression variation, including the prominent effects of local expression quantitative trait loci (eQTL), where a gene’s mRNA abundance is strongly associated with genotype near its genomic location. There are also *trans-* acting effects, where single loci affect the mRNA abundance of large numbers of genes located throughout the genome.

To identify genes and pathways that contribute to obesity-induced type II diabetes, we constructed a large mouse F_2_ intercross between diabetes-resistant (C57BL/6J, abbreviated B6) and diabetes-susceptible (BTBR *T*^+^ *tf* /J, abbreviated BTBR) mouse strains. Greater than 500 F_2_ offspring were generated. All were genetically obese through introgression of the leptin mutation (*Lep*^*ob/ob*^) and all were sacrificed at 10 weeks of age, the age when essentially all BTBR ob/ob mice are diabetic. Thus, the screen focused on genetic differences between the B6 and BTBR mouse strains. In addition to measuring numerous diabetes-related clinical phenotypes (e.g., circulating levels of insulin and glucose), we conducted genome-wide gene expression profiling in six tissues of every F_2_ mouse; adipose, gastrocnemius muscle, hypothalamus, pancreatic islets, kidney, and liver.

We identified numerous *trans*-eQTL hotspots. Some were common to more than one tissue, whereas others were strongly tissue-specific. We identified a particularly striking *trans*-eQTL hotspot on distal chromosome 6, at ∼142 Mb. This hotspot was observed exclusively in pancreatic islets and affected ∼2,400 transcripts encoded by genes located throughout the genome. Using principal component analysis of the transcripts mapping to the locus with the strongest LOD scores (LOD ∼100), we were able to infer the eQTL genotype for all mice, thereby converting the multivariate gene expression phenotype to a co-dominant Mendelian trait. We applied the traditional method for fine-mapping a Mendelian trait. It involved identifying recombinant mice and genotyping additional markers to precisely define the location of the recombination events. This allowed us to refine the location of the *trans*-eQTL to a 298 kb interval containing just three genes.

Among the three genes included in the 298kb region are two members of the *SLCO* gene family, *Slco1a5* and *Slco1a6*, which encode the organic anion transporting polypeptide (OATP) 1A5 and OATP1A6 (Hagenbuch and Stieger, 2013). OATPs have been shown to transport numerous endogenous substrates, including bile acids such as taurocholic acid (Hagenbuch and Stieger, 2013). In addition to their role as detergents, bile acids have hormone-like properties due to their ability to activate nuclear hormone receptors (Parks et al. 1999). This makes genetic variation in a bile acid transporter a plausible mechanism for the differential regulation of numerous genes. We characterized functional differences between the two *Slco1a6* variants of B6 and BTBR mice and conclude that genetic variation in *Slco1a6* leads to altered transport of taurocholic acid (TCA), and potentially other bile acids, by pancreatic islets, resulting in broad gene regulation.

## Materials and Methods

### Mice and genotyping

C57BL/6J (abbreviated B6 or B) and BTBR *T*^+^ *tf* /J (abbreviated BTBR or R) mice were purchased from the Jackson Laboratory (Bar Harbor, ME) and bred at the Biochemistry Department at the University of Wisconsin–Madison. The *Lep*^*ob/ob*^ mutation, which originated in the B6 strain (Ingalls et al. 1950), was introgressed into the BTBR strain. F_1_ *Lep*^*ob/ob*^ were made fertile by employing adipose tissue transplants from wild type mice to restore leptin.

All mice were genotyped with the 5K GeneChip (Affymetrix). A large set of DNA sample mix-ups were identified, by comparing observed genotypes to predictions based on large-effect local eQTL, and corrected (Broman *et al.* 2014). After data cleaning, there were 519 F_2_ mice genotyped at 2,057 informative markers, including 20 on the X chromosome. The leptin gene resides on proximal chromosome 6, at ∼29.0 Mbp, and the proximal 32 Mbp of this chromosome showed marked segregation distortion, with excess B6 homozygotes and reduced BTBR homoyzogotes. However, the *trans*-eQTL hotspot under study is at the opposite end of the chromosome from the leptin gene, and the region around the hotspot segregated normally.

### Gene expression microarrays

Gene expression was assayed with custom two-color ink-jet microarrays manufactured by Agilent Technologies (Palo Alto, CA). RNA preparations were performed at Rosetta Inpharmatics (Merck & Co.). Six tissues from each F_2_ mouse were used for expression profiling; adipose, gastrocnemius muscle (abbreviated gastroc), hypothalamus (abbreviated hypo), pancreatic islets (abbreviated islet), kidney, and liver. Tissue-specific mRNA pools for each tissue were used for the reference channel, and gene expression was quantified as the ratio of the mean log_10_ intensity (mlratio). For further details, see Keller *et al.* (2008). In the final data set, there were 519 mice with gene expression data on at least one tissue (487 for adipose, 490 for gastroc, 369 for hypo, 491 for islet, 474 for kidney, and 483 for liver). The microarray included 40,572 total probes; we focused on the 37,797 probes with known location on one of the autosomes or the X chromosome.

### QTL analysis

For QTL analysis, we first transformed the gene expression measures for each microarray probe in each of the six tissues to normal quantiles, taking Φ^-1^[(*R*_*i*_ – 0.5)/*n*], where Φ is the cumulative distribution function for the standard normal distribution and *R*_*i*_ is the rank in {1, …, *n*} for mouse *i*. We then performed single-QTL genome scans separately for each probe in each tissue, by Haley-Knott regression (Haley and Knott 1992) with microarray batch included as an additive covariate and with sex included as an interactive covariate (i.e. allowing the effects of QTL to be different in the two sexes). Calculations were performed at the genetic markers and at a set of pseudomarkers inserted into marker intervals, selected so that adjacent positions were separated by ≤ 0.5 cM. We calculated conditional genotype probabilities, given observed multipoint marker genotype data, using a hidden Markov model assuming a genotyping error rate of 0.2%, and with genetic distances converted to recombination fractions with the Carter-Falconer map function (Carter and Falconer 1951).

For each probe in each tissue, we focused on the single largest LOD score peak on each chromosome, and LOD score peaks ≥ 5 (corresponding to genome-wide significance at the 5% level, for a single probe in a single tissue, determined by computer simulations under the null hypothesis of no QTL).

### Inference of eQTL genotype

To infer the eQTL genotype of each mouse at the islet chromosome 6 *trans-*eQTL hotspot, we selected the 181 microarray probes that mapped to the locus with LOD ≥ 100, but were not associated with a gene located anywhere on chromosome 6, thereby ensuring they represented a *trans*-eQTL. We computed the first two principal components, using the mlratio measure of gene expression (i.e. prior to the normal quantile transformation that was used in the QTL analyses). The scatterplot of the two principal components revealed three distinct clusters of mice, corresponding to BB (homozygous B6), BR (heterozygous) or RR (homozygous BTBR). We used the mice without a recombination event in the region to define the genotype for these clusters, from which we could then infer the eQTL genotype for the mice with a recombination event in the region of the QTL.

### Fine-mapping the eQTL

Fine-mapping of the chromosome 6 islet eQTL proceeded by the traditional method for a simple Mendelian trait: we identified the smallest interval in which individuals’ genotypes at genetic markers were consistent with their inferred eQTL genotype. We selected individuals with recombination events flanking this interval and genotyped these recombinant animals at additional SNP markers in the interval, in order to more precisely localize their recombination events and so refine the QTL interval. We used SNPs predicted from high-throughput sequencing of genomic DNA from two BTBR mice (Eric Schadt, Mt. Sinai, personal communication) to guide our initial selection of the additional markers that were used to genotype recombinant individuals. Predicted SNPs proximal to the eQTL locus on chromosome 6 were confirmed with PCR amplification followed by Sanger Sequencing in BTBR and B6 genomic DNA. Confirmed SNPs were then used to further map the eQTL region using F_2_ mice whose recombinations were within the eQTL interval. File S1 provides a complete list of the SNPs that were used to narrow our eQTL interval.

### Characterization of the transport protein OATP1A6 encoded by *Slco1a6* gene

Radiolabeled [^3^H]-taurocholic acid (10 Ci/mmol), [^3^H]-cholic acid (30 Ci/mmol) and [^3^H]- methotrexate (21.6 Ci/mmol) were purchased from American Radiolabeled Chemicals, Inc. Radiolabeled [^3^H]-estrone-3-sulfate (45 Ci/mmol), [^3^H]- (D-Pen^2^, D-Pen^5^)-enkephalin (44 Ci/mmol), [^3^H]-estradiol-17-β-glucuronide (41.4 Ci/mmol) were obtained from PerkinElmer (Boston, MA). [^3^H]-bromosulfophthalein (11.5 Ci/mmol) was obtained from International Isotope Clearing House (Leawood, KS, USA). All other chemicals used for the characterization were purchased from Sigma-Aldrich.

### Cloning of mouse OATP1A6 and expression in HEK293 cells

The open reading frame (ORF) encoding mouse OATP1B6 was PCR amplified and cloned using cDNA prepared from RNA that was isolated from kidneys from B6 or BTBR mice using the following primers: forward primers containing an *NheI* restriction site were: 5’- AGAGGCTAGCACCATGGGAGAACCTGGGA-3’ for B6, and 5’-AGAGGCTAGCACCATGGGAGAACCTGAGA-3’ for BTBR, and a reverse primer for both strains containing a *Not I* restriction site: 5’- AGAGGCGGCCGCCCTACAGCTTAGTTTTCAGTTCTCCA-3’. The PCR products were first cloned directionally into pcDNA5/FRT. In order to achieve the same level of expression for the B6 *vs.* BTBR ORFs, they were subcloned into a vector that contained the 3’untranslated region of mouse OATP1A6 (strain FVB/N, accession number BC071214; purchased from Thermo Fisher Scientific, formerly Open Biosystems). All clones were sequence verified on both strains. Human embryonic kidney (HEK) 293 cells (ATCC, Manassas, VA) were grown at 37 °C in a humidified 5% CO_2_ atmosphere in DMEM High Glucose (ATCC) supplemented with 10% FBS (Hyclone), 100 U/ml penicillin, and 100 μg/ml streptomycin. HEK293 cells were plated at 250,000 cells per well in 24-well-plates pre-coated with 0.1 mg/ml poly-D-lysine. Twenty-four hours later cells were transfected with 0.5 μg plasmid DNA and 1.5 ul Fugene HD (Promega) per well. The uptake assays described below were performed 48 hours following transfection.

### Site directed mutagenesis of mouse OATP1A6

The serine at position 564 in B6 was mutated to a proline and the proline at position 564 in BTBP to a serine using the Quick Change site directed mutagenesis kit (Agilent Technologies, Santa Clara, CA) using the following primers: 5’- CAAAGCGCCAAAGTAAATAGATGCAGGAATACCAGCAAATA-3’ and 5’- TATTTGCTGGTATTCCTGCATCTATTTACTTTGGCGCTTTG-3’ for B6; 5’- CAAAGCGCCAAAGTAAATAGGTGCAGGAATACCAGCAAATA-3’ and 5’- TATTTGCTGGTATTCCTGCACCTATTTACTTTGGCGCTTTG-3’ for BTBR. All mutants were verified by DNA sequencing.

### Uptake activity assays for mouse OATP1A6

HEK293 cells transfected with either an empty vector (EV), the ORF encoding B6 OATP1A6 or the ORF encoding BTBR OATP1A6 were washed three times with 1 ml of pre-warmed (37°C) uptake buffer containing 142 mM NaCl, 5 mM KCl, 1 mM KH_2_PO_4_, 1.2 mM MgSO_4_, 1.5 mM CaCl_2_, 5 mM glucose, and 12.5 mM HEPES, pH 7.4. Following the wash, 200 μl of uptake buffer (37 °C) containing the radiolabeled substrates were added to the well. At indicated times, uptake was terminated by four washes of 1 ml of ice-cold uptake buffer. Cells were then lysed with 300 μl 1% Triton X-100 in PBS at room temperature for 20 min. Two hundred μl of cell lysate were transferred to a 24-well scintillation plate (Perkin Billerica, MA) and 750 μl Optiphase Supermix scintillation cocktail (Perkin Elmer) was added to each well. Radioactivity was measured in a Microbeta Trilux liquid scintillation counter. The remaining cell lysates were transferred to 96-well plates to determine total protein concentration using the bicinchoninic acid protein assay (Pierce Biotechnology, Inc. Rockford, IL). All transport measurements were corrected by the total protein concentration and by the specific activities of the different substrates to yield pmoles per mg protein. All experiments were performed three independent times, with each study including triplicate technical measurements.

### Software

All statistical analyses were conducted with R (R Development Core Team 2015). QTL analyses were performed with the R package, R/qtl (Broman *et al.* 2003).

### Data availability

The genotype and gene expression microarray data are available at the QTL Archive, now part of the Mouse Phenome Database: http://phenome.jax.org/db/q?rtn=projects/projdet&reqprojid=532

## Results

### Identification of eQTL

We performed a single-QTL genome scan for all genes in six tissues of ∼500 F_2_ mice constructed from an intercross of diabetes-resistant (B6) and diabetes-susceptible (BTBR) mice. For all genes, we identified the single largest LOD score on each chromosome. The inferred expression quantitative trait locus (eQTL) for all genes with LOD ≥ 5 are displayed in Figure 1, with the y-axis corresponding to the genomic position of the microarray probe and the x-axis corresponding to the estimated eQTL position. As expected, we see a large number of local-eQTL along the diagonal for each tissue-specific panel. These local-eQTL correspond to genes for which expression or mRNA abundance is strongly associated with genotype near their genomic position and are often referred to as cis-eQTL.

**Figure 1.**
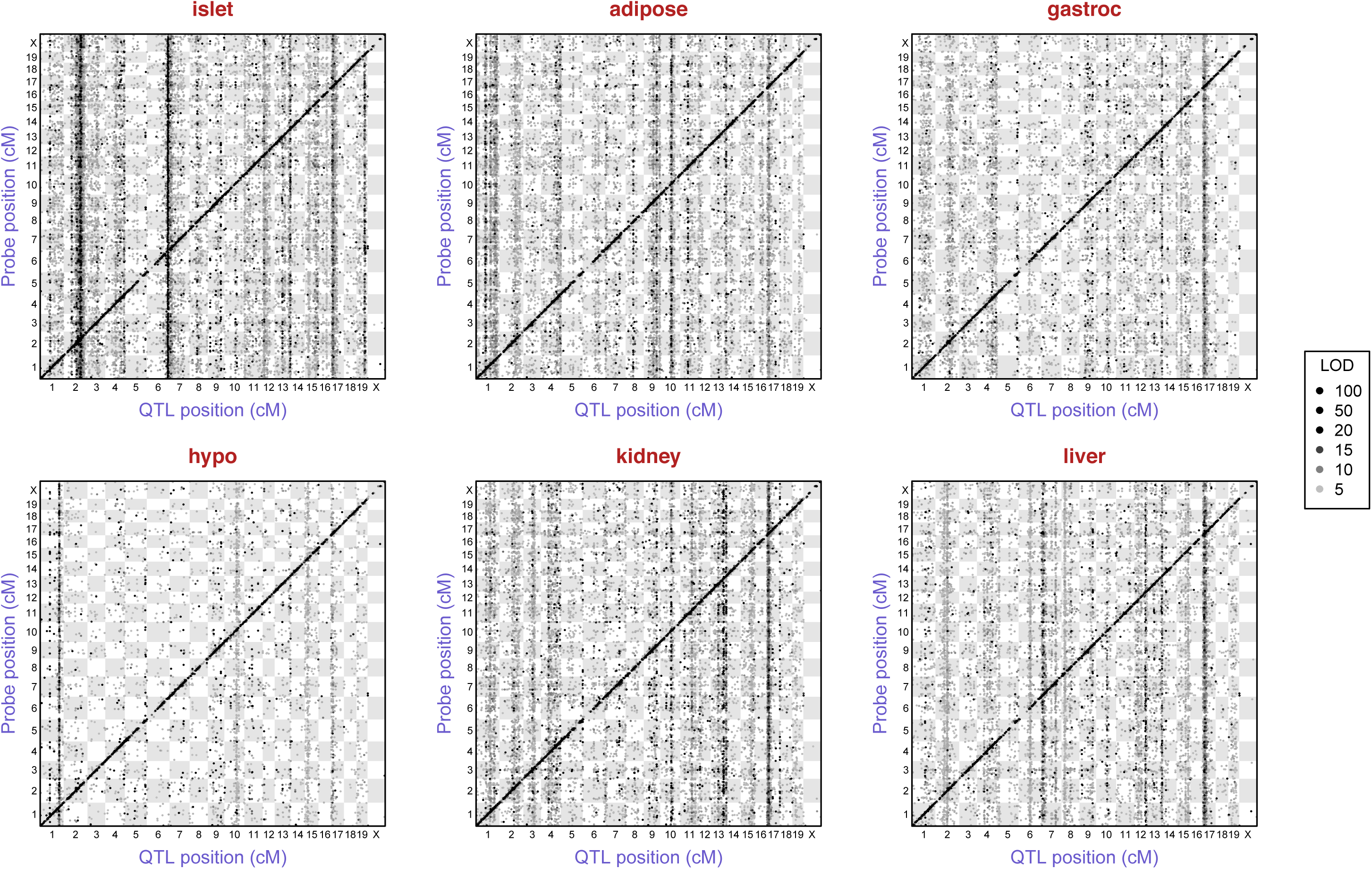
Inferred eQTL with LOD ≥ 5, by tissue. Points correspond to peak LOD scores from single-QTL genome scans with each microarray probe with known genomic position. The y-axis is the position of the probe and the x-axis is the inferred QTL position. Points are shaded according to the corresponding LOD score, though we threshold at 100: all points with LOD ≥ 100 are black.

In addition to the local-eQTL, there are a number of prominent vertical bands: genomic loci that influence the expression of genes located throughout the genome. These distally-mapping eQTL are often referred to as *trans*-eQTL, and by virtue of their common genetic architecture (co-mapping), yield the *trans*-eQTL hotspots.

Overall, we detected many more *trans*-eQTL than *cis*-eQTL. The *trans*-eQTL hotspots can show either remarkable tissue specificity or be observed in multiple tissues. For example, a locus near the centromere of chromosome 17 (11.73 cM) shows effects in all tissues, with an average of 1498 microarray probes mapping to this region with LOD ≥ 5 in the tissues surveyed. In contrast, a *trans*-eQTL hotspot located at the distal end of chromosome 6 was only observed in pancreatic islets. Furthermore, this islet-specific *trans*-eQTL hotspot was the strongest that we detected, with more eQTL mapping to this locus than any other locus identified in our study. We chose this *trans*-eQTL hotspot for further study.

Figure 2 displays, for each tissue, the estimated QTL location and LOD score for all probes mapping to chromosome 6 with LOD ≥ 5. The peak marker for the chromosome 6 *trans*-eQTL hotspot is rs8262456, located at 91.4 cM (141.52 Mbp in NCBI build 37). For pancreatic islets, in the 10 cM interval centered at this marker, there are 2889 probes with LOD peaks ≥ 5 (7.6% of the 37,797 probes considered), including 1700 probes with LOD ≥ 10 (4.5% of all probes) and 199 probes with LOD ≥ 100. These 2889 co-mapping probes are located throughout the genome, including all autosomes and the X chromosome, and include 756 with no annotated gene and 2133 probes with a gene annotation, corresponding to 2085 distinct genes (2038 genes with one probe, 46 genes with two probes, and one gene with three probes). This *trans*-eQTL hotspot is specific to islet cells; the other five tissues have 66–140 probes mapping to this 10 cM interval with LOD ≥ 5, and 36–66 probes with LOD ≥ 10. These results suggest that a gene located at ∼141.5 Mbp on chromosome 6 regulates the expression of genes throughout the genome in an islet-specific fashion.

**Figure 2.**
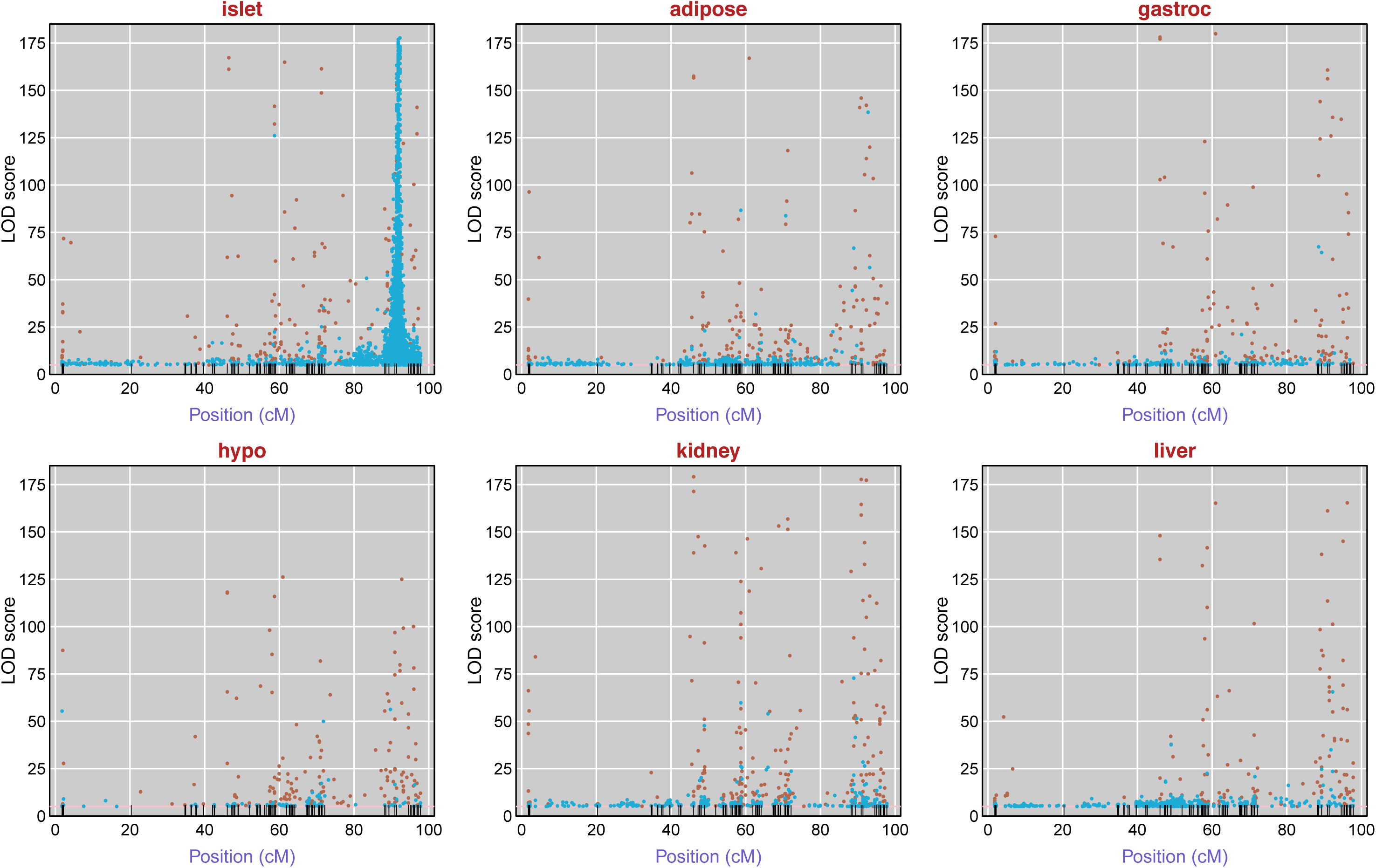
Inferred chromosome 6 eQTL with LOD ≥ 5, by tissue. Each point corresponds to a microarray probe, and indicates the maximum LOD score on chromosome 6 for that probe, and the position of the peak LOD score. Brown points correspond to probes whose genomic location is on chromosome 6; blue points are for probes on other chromosomes.

The *trans*-eQTL shows approximately additive allele effects in all microarray probes (Figure S1), and approximately equal numbers of probes having an effect in each direction: Among the 2889 probes mapping with LOD ≥ 5 to the 10 cM interval centered at the peak marker for the *trans*-eQTL, there are 1432 for which the BTBR homozygote has higher expression than the B6 homozygote, and 1457 for which the BTBR homozygote has lower expression.

### Prediction of QTL genotype from phenotype

To identify gene candidates at the islet chromosome 6 locus, our first step was to narrow the effective interval. We calculated the first two principal components of the expression values for the 181 microarray probes that mapped with LOD ≥ 100 to the 10 cM interval centered at the peak marker for the chromosome 6 islet *trans-*eQTL hotspot but that did not reside on chromosome 6. (We considered probes with local-eQTL to be more likely affected by a separate polymorphism.) A scatterplot of these two principal components reveals three distinct clusters of mice (Figure 3). Points corresponding to mice without a recombination event in the 10 cM interval are colored according to their genotype in the interval. This indicates that the three clusters correspond to the three possible eQTL genotypes. There were 74 mice with recombination events in the interval (displayed in yellow in Figure 3). We inferred the eQTL genotype for these recombinant mice, based on the cluster in which they reside. In this manner, the multivariate gene expression phenotype was converted to a co-dominant Mendelian phenotype.

**Figure 3.**
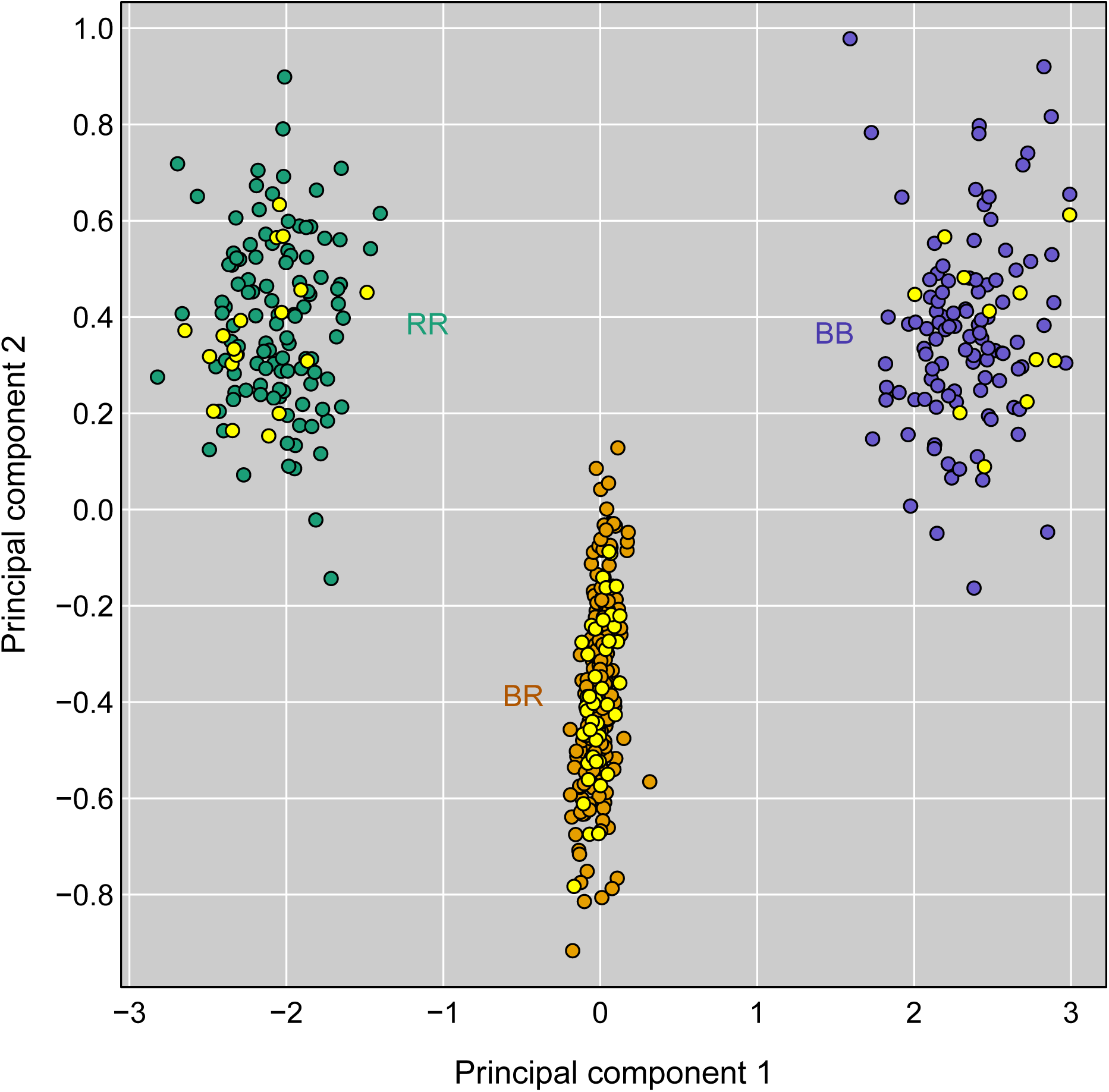
The first two principal components for the islet gene expression data for the 181 microarray probes that map to the chromosome 6 *trans*-eQTL hotspot with LOD ≥ 100 but do not reside on chromosome 6. Each point is a mouse; points for mice without a recombination event in the 10 cM interval centered at the peak marker are colored by their genotype in the region. Yellow points correspond to the 74 mice with recombination events in the interval.

### Fine-mapping the eQTL

By comparing the inferred eQTL genotypes at the islet chromosome 6 *trans-*eQTL hotspot to the observed marker genotypes in the region, we localized the QTL to the 3.4 Mbp interval between markers rs8262456, at 141.52 Mbp, and rs13479085, at 144.91 Mbp; see Figure 4A, in which the diamonds represent the inferred eQTL genotypes. There were 29 mice with recombination events in this interval.

**Figure 4.**
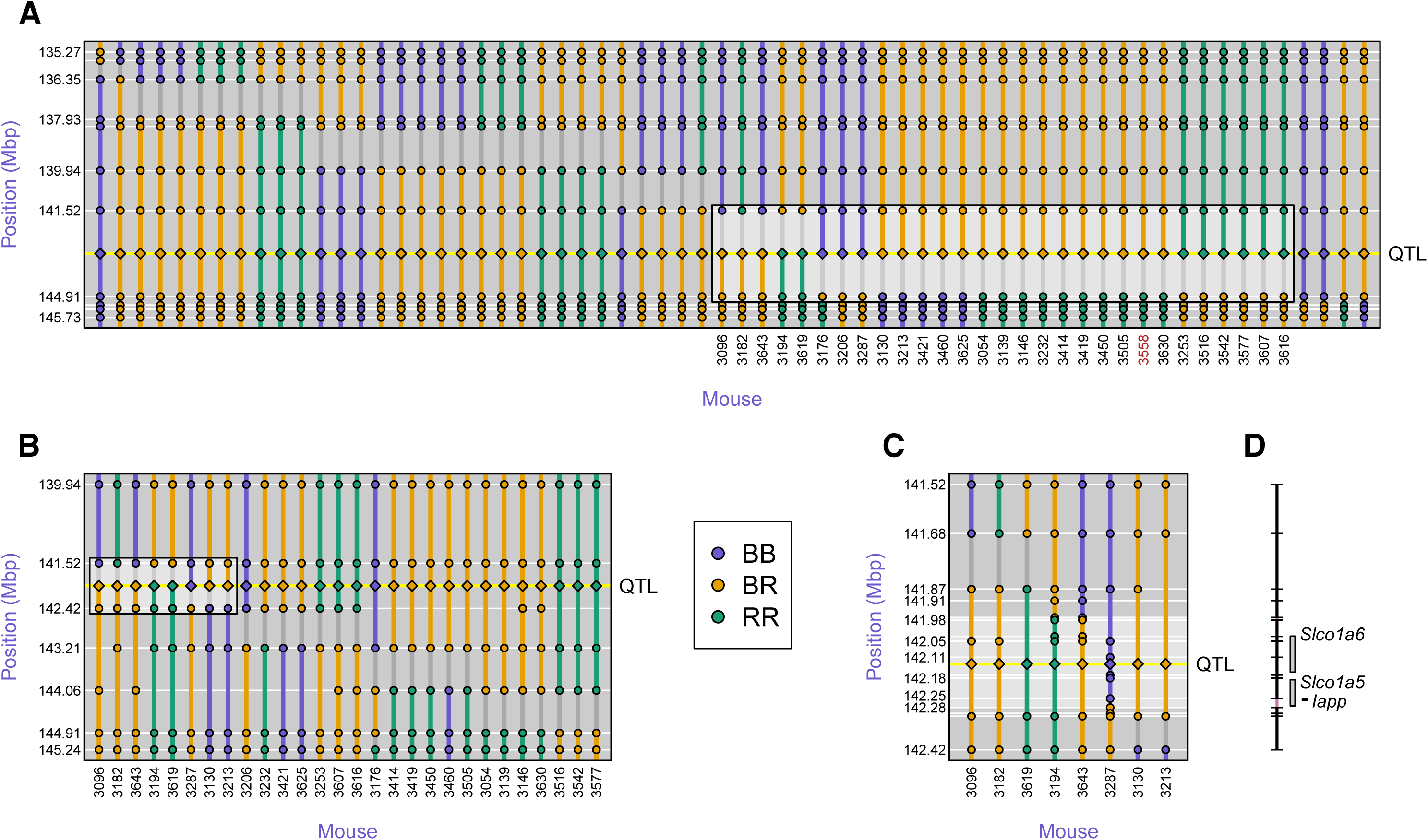
Fine-mapping of the islet chromosome 6 eQTL. **A:** Initial SNP genotypes of the 64 mice with recombination events in the 10 Mbp (7 cM) region around the QTL, along with their inferred QTL genotypes (shown at the center of the inferred interval). The highlighted box indicates 29 mice with recombination events in the QTL interval. **B:** Additional genotypes on five markers in the QTL interval, for 28 of the 29 mice with recombination event flanking the QTL. The highlighted box indicates eight mice with recombination event flanking the QTL. **C:** Additional genotypes in the reduced QTL interval, for eight mice with recombination events flanking the QTL. The QTL interval is further reduced to 298 kbp (141.979 – 142.277 Mbp), a region containing three genes (**D**).

Genotyping of 5 additional markers in the region (Figure 4B) refined the QTL position to a 904 kb interval, from 141.517 – 142.421 Mbp. There were eight mice with recombination events in this interval.

Genotypes at an additional 14 markers in these eight mice (Figure 4C) refined the QTL position to the 298 kb interval from 141.979 – 142.277 Mbp. The interval contains three genes, *Slco1a6*, *Slco1a5*, and *Iapp*. Recombination events in three mice define this interval: two at the proximal end and one at the distal end. Additional genotyping that refined the locations of these recombination events would not permit us to exclude any of the three genes. The expression levels of *Slco1a6* and *Iapp* are strongly associated with genotype in the region, with LOD scores of 161 and 10.6, respectively. *Slco1a5*, on the other hand, has a maximum LOD score on chromosome 6 of 2.4 (and at the other end of the chromosome) and shows evidence for a QTL on chromosome 7, with a LOD score of 9.2. That *Slco1a6* shows such a strong local-eQTL makes it our strongest candidate.

### Characterization of *Slco1a6*

To characterize the genetic differences between B6 *vs.* BTBR *Slco1a6*, we sequenced all the exons from the *Slco1a6* gene in both mouse strains. In addition, we cloned cDNA prepared from total kidney RNA purified from each strain. Compared to B6 as the reference sequence, the BTBR sequence differed in 9 nucleotides, resulting in 6 amino acid changes (Table S1). Alignment of the amino acid sequence for BTBR OATP1A6 (the gene product for mouse *Slco1a6*) with 10 other OATP1A family members from human, rat and mouse (including the B6 reference sequence), revealed the degree of conservation for these 6 amino acid changes. Two SNPs were selective for either B6 or BTBR among the OATP1A sub-family: 1) at position 5, B6 OATP1A6 is a Glycine residue, whereas in all other proteins of this sub-family this residue is a Glutamate residue; and 2) at position 564, BTBR OATP1A6 is a Serine residue, whereas it is a Proline residue in the other family members. At amino acid positions 456 and 658, B6 OATP1A6 and the human OATP1A2 are the same, and different than all other members of the sub-family as well as BTBR OATP1A6. Finally, the remaining two differences between B6 and BTBR OATP1A6 were also different among the other family members.

Molecular modeling of the protein structure for mouse OATP1A6 suggests that the conversion of the highly conserved Pro^564^ forms a kink in the alpha-helix for one of the trans-membrane domains, and this Pro residue faces into the central aqueous pore of the protein through which substrates are translocated (Figure 5A) (Meier-Abt *et al.* 2005). Conversion of this to Ser^564^ in BTBR OATP1A6 may alter the substrate selectivity and/or the transport activity of the protein. The Gly^5^ difference between B6 and BTBR OATP1A6 falls into the unstructured amino-terminal region of the protein.

**Figure 5.**
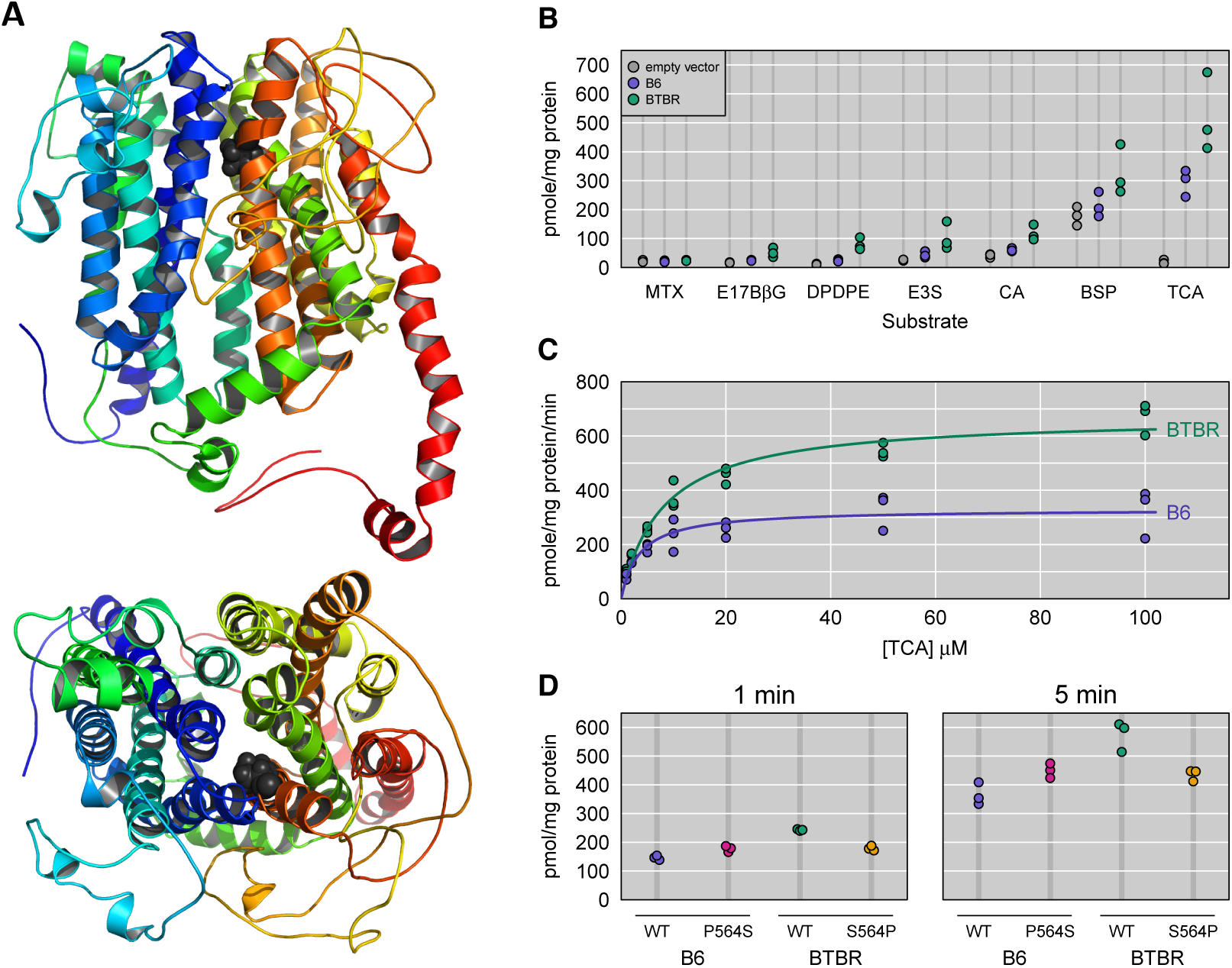
Functional characterization of mouse OATP1A6. **A:** Pymol views of a homology model of mouse OATP1A6 with the proline at position 564 highlighted **B:** Mouse OATP1A6-mediated uptake of some common OATP substrates was determined in HEK293 cells transiently transfected with empty vector (EV, gray points), C57BL/6-OATP1A6 (blue points) or BTBR-OATP1A6 (green points). Uptake of 10 μM taurocholic acid (TCA), cholic acid (CA), estrone-3-sulfate (E3S), methotrexate (MTX), (D-Pen^2^, D-Pen^5^)-enkephalin (DPDPE), estradiol-17-*β*-glucuronide (E17*β*G), and bromosulfophthalein (BSP) was measured at 37 °C for 5 min. **C:** Uptake of TCA was measured at various concentrations from 1 to 100 μM at 37 °C for 1 min with empty vector and B6- or BTBR-OATP1A6- expressing HEK293 cells. Net uptake was calculated by subtracting the values of empty vector-transfected cells from either B6- or BTBR-OATP1A6 transfected cells. Resulting data were fitted to the Michaelis-Menten equation to obtain *K*_m_ and *V*_max_ values. **D:** Uptake of 10μM TCA was measured at 1 and 5 min with normal B6 OATP1A6-Pro^564^ or mutant Ser^564^ and with normal BTBR OATP1A6-Ser^564^ or mutant Pro^564^. For the uptake experiments, each point shown is derived from an individual experiment performed with triplicate determinations.

To functionally characterize mouse OATP1A6, encoded by *Slco1a6*, and to evaluate potential differences between the B6 and BTBR proteins that may be related to the coding variants we detected in our sequencing, we performed cellular uptake studies of several known substrates for the OATP family of transporters. HEK293 cells were transiently transfected with the complete open reading frame (ORF) encoding either B6 or BTBR OATP1A6. With the exception of methotrexate (MTX), the uptake of all other substrates was significantly enhanced in cells overexpressing either variant of mouse OATP1A6 (Figure 5B). Furthermore, cells expressing BTBR OATP1A6 showed significantly elevated uptake of several substrates compared to cells expressing B6 OATP1A6. Of all substrates tested, uptake of the primary conjugated bile acid, taurocholate (TCA) showed the greatest signal above empty-vector control cells, for both B6 and BTBR OATP1A6. The primary unconjugated bile acid cholate (CA) was also transported, but less efficiently than TCA (Figure 5B) whereas taurochenodeoxycholic acid (TCDCA) was not a substrate for either B6 or BTBR OATP1A6 (data not shown). These results suggest that mouse OATP1A6 can mediate cellular uptake of certain bile acids and that one or more of the polymorphisms we identified in B6 vs. BTBR OATP1A6 yielded enhanced transport activity for the BTBR protein.

The studies illustrated in Figure 5B surveyed the substrate selectivity of mouse OATP1A6. In order to evaluate kinetic differences between B6 and BTBR OATP1A6, we measured the TCA uptake at increasing concentrations in HEK cells overexpressing one or the other mouse variant (Figure 5C). Uptake measurements were made at 1 min (initial linear range: 0 to 2 min at low concentration and 0 to 1 min at high concentration). BTBR-OATP1A6 has a much higher transport capacity (V_max_ = 673 ± 22 pmol/mg protein x min) than B6-OATP1A6 (V_max_ = 330 ± 18 pmol/mg protein x min). The calculated affinity for TCA is slightly lower for BTBR OATP1A6 (K_m_ = 8 ± 1μM) than for B6 OATP1A6 (K_m_ = 3.5 ± 0.8 μM). Because position 564 could be functionally important, we mutated this position in the two OATPs to the corresponding amino acid found in the other protein and tested them. As compared to B6- OATP1A6, B6-OATP1A6 with a serine at position 564 showed a 1.2 fold increased TCA uptake (Figure 5D). Similarly, as compared to BTBR-OATP1A6, BTBR-OATP1A6, with a proline at position 564, showed a 1.3 fold decreased TCA uptake (Figure 5D). These results support the hypothesis that position 564 is crucial for OATP1A6 function and/or expression.

## Discussion

Analysis of gene expression variation in a large mouse intercross revealed a *trans*-eQTL hotspot on distal chromosome 6 (Figures 1 and 2), affecting the mRNA abundance for ∼8% of the ∼22,000 annotated genes, many with LOD scores ≥ 100. This *trans*-eQTL hotspot affected more genes than any other hotspot detected in the six tissues profiled, and was solely observed in pancreatic islets, adding to its unique character. Principal component analysis of the *trans*-eQTL hotspot revealed three distinct clusters of mice (Figure 3) that corresponded to the three eQTL genotypes. This allowed us to infer the eQTL genotypes of all mice and, in doing so, convert the multivariate gene expression phenotype to a co-dominant Mendelian trait. With the initial marker genotypes, the eQTL could be localized to a 3.4 Mbp interval. Following two rounds of genotyping with additional markers in recombinant mice (Figure 4), we were able to reduce this to a 298 kb interval containing just three genes: *Slco1a5*, *IAPP*, and *Slco1a6*.

Only *Slco1a6* mRNA showed a clearly different expression pattern between B6 and BTBR mice, and it showed significantly higher expression than *Slco1a5* mRNA (Keller *et al.* 2008), and so we focused on *Slco1a6*. OATP1A6, the protein encoded by mouse *Slco1a6*, was reported to be a kidney-specific organic anion transporter whose mRNA was not detectable for the first three weeks after birth in BALB/c mice and reached adult levels more than six weeks after birth (Choudhuri *et al.* 2001), but no functional data has been published. Because most OATPs are able to transport bile acids, we first tested whether OATP1A6 was able to mediate uptake of bile acids. Indeed, we demonstrated that both conjugated (TCA) as well as unconjugated bile acids (CA) are substrates for OATP1A6 (Figure 5B). In addition, like many other OATPs (Roth *et al.* 2012), OATP1A6 is a multispecific transporter and can transport not just bile acids but also the hormone conjugates estradiol-17β-glucuronide and estrone-3-sulfate, the dye bromosulfophthalein and the opioid peptide (D-Pen^2^, D-Pen^5^)-enkephalin (DPDPE). All the tested substrates were consistently better transported by BTBR OATP1A6 as compared to B6 OATP1A6 (Figure 5B,C). Among the 6 amino acid changes in OATP1A6, between B6 and BTBR, position 564 is the most likely to be involved in the functional difference between the two OATPs. The functional data with the mutants confirm this suggestion, at least in part, and demonstrate that indeed the substitution of the proline to serine at position 564 in B6 results in a transporter that shows increased TCA uptake (Figure 5D), and vice versa; that substitution of serine at position 564 in BTBR to proline decreases TCA transport. Although these single amino acid substitutions do not convert the B6 into the BTBR protein, the functional trend supports the suggested importance of this amino acid position. Additional experiments are required to determine whether these changes affect protein expression levels or turnover numbers.

Abe *et al.* (2010), using immunohistochemistry, demonstrated that the closely related OATP1A1 was expressed in β-cells and that OATP1A5 was expressed in islets and acinar cells of rat pancreas. In contrast, OATP1A4 was shown to be expressed in α-cells (Abe *et al.* 2010).

However, the antibody used to detect rat OATP1A1 in the pancreas was raised against the C-terminal end of the protein, which is identical in OATP1A1 and OATP1A6, and therefore this antibody is not able to distinguish between these two OATPs. Given that obesity-dependent upregulation in BTBR mice was demonstrated for *Slco1a6* but not *Slco1a1* mRNA (Keller *et al.* 2008), it is unlikely, at least in mouse pancreas, that OATP1A1 is expressed. The suggested OATP1A1-mediated pravastatin uptake into β-cells to increase insulin secretion (Abe *et al.* 2010) could also be explained by OATP1A6-mediated pravastatin transport (data not shown).

An OATP1A6 knockout mouse has so far not been reported. However, OATP1A/1B knockout mice where the whole *Slco1a/1b* locus was deleted are viable and have increased plasma levels of unconjugated bile acids as well as bilirubin and bilirubin monoglucuronide levels (van de Steeg *et al.* 2010) further demonstrating that bile acids are important substrates of OATPs.

While we have not yet tested the effect of bile acids on the transcripts that map to the chromosome 6 *trans-*eQTL hotspot, bile acids can affect gene regulation via farnesoid X receptor (FXR), a nuclear receptor and transcription factor that is activated by bile acids. Moreover, Düfer et al. (2012) showed that certain bile acids can stimulate insulin secretion via FXR activation.

There has been some controversy about *trans*-eQTL hotspots potentially being artifacts (for example, see Breitling *et al.* 2008; Kang *et al.* 2008). If multiple transcripts show strongly correlated expression, perhaps induced by some confounding factor, such as a batch effect, and if one such transcript shows chance association to genotype at some location, then many transcripts will share this common, false-positive eQTL. This has led to the development of a number of methods to control for underlying confounding factors (Leek and Storey 2007; Kang *et al.* 2008; Stegle *et al.* 2010; Listgarten *et al.* 2010; Gagnon-Bartsch and Speed 2012; Fusi *et al.* 2012). However, the very strong associations to our chromosome 6 *trans*-eQTL hotspot (with 177 transcripts, from other chromosomes, mapping to the region with LOD ≥ 100), precludes the possibility that this is an artifact due to confounding: If these associations arose due to some underlying confounder, such as a batch effect, the strength of the association between the confounder and genotype in the region would have to be yet larger.

Our approach to convert the multivariate gene expression phenotype, for transcripts mapping to the chromosome 6 *trans*-eQTL hotspot, to a simple Mendelian trait could be applied more generally. While it was sufficient for us to consider the first two principal components to define the clusters of mice with a common eQTL genotype, one could focus on the non-recombinant mice (whose eQTL genotype is known) and apply linear discriminant analysis or other classification algorithms in order to infer the eQTL genotype in the recombinant mice. A key consideration that must be addressed is the possibility of multiple linked polymorphisms.

Efforts to improve the QTL mapping precision in experimental crosses have focused on increasing the density of recombination events, e.g. with advanced intercross lines (Darvasi and Soller 1995) and heterogeneous stock (Mott *et al.* 2000). But the precision of QTL mapping may be hampered more by residual phenotype variation than by lack of recombination events. With a co-dominant Mendelian trait in an intercross, the distribution of the length of the interval defined by recombination events flanking the trait locus follows a gamma distribution with shape=2 and rate=2*n*, where *n* is the sample size. With 100 intercross mice, the average interval is just 1 cM. This points to the importance of integrating multiple phenotypes that map to a common locus in order to develop composite traits with markedly reduced residual variation.

## Acknowledgments

The authors thank Amit Kulkarni for providing annotation information for the gene expression microarrays, Eric Schadt for providing predicted SNPs based on his high-throughput sequencing of two BTBR mice, and Petr Simecek for his assistance in preparing the data for the Mouse Phenome Database. This work was supported in part by National Science Foundation grant DMS-12-65203 (to C.K.) and National Institutes of Health grants GM074244 (to K.W.B.), DK066369 (to A.D.A), GM077336 (to B.H.), and GM102756 (to C.K.)

